# On Binocular Vision: The Geometric Horopter and Cyclopean Eye

**DOI:** 10.1101/037069

**Authors:** Jacek Turski

**Affiliations:** Department of Mathematics and Statistics^1^ University of Houston-Downtown

**Keywords:** Binocular vision, Horopter, Cyclopean eye, Disparity

## Abstract

We study geometric properties of horopters defined by the criterion of equality of angle. Our primary goal is to derive the precise geometry for anatomically correct horopters. When eyes fixate on points along a curve in the horizontal visual plane for which the vergence remains constant, this curve is the larger arc of a circle connecting the eyes’ rotation centers. This isovergence circle is known as the Vieth-Müller circle. We show that, along the isovergence circular arc, there is an infinite family of horizontal horopters formed by circular arcs connecting the nodal points. These horopters intersect at the point of symmetric convergence. We prove that the family of 3D geometric horopters consists of two perpendicular components. The first component consists of the horizontal horopters parametrized by vergence, the point of the isovergence circle, and the choice of the nodal point location. The second component is formed by straight lines parametrized by vergence. Each of these straight lines is perpendicular to the visual plane and passes through the point of symmetric convergence. Finally, we evaluate the difference between the geometric horopter and the Vieth-Müller circle for typical near fixation distances and discuss its possible significance for depth discrimination and other related functions of vision that make use of disparity processing.

## 1 Introduction

In the primate visual system, basic concepts of binocular projection include the horopter and the Cyclopean eye. The horopter is the locus of points in space seen singly by the two eyes. The Cyclopean eye is an abstract eye that represents the visual axes of the two eyes by a single axis of perceived direction.

An eye model that still influences theoretical developments in binocular vision assumes that the optical nodal point coincides with the center of rotation for eye movements. This anatomically incorrect assumption was originally made about two centuries ago in the construction of Müller’s horopter, known as the Vieth-Müller circle (V-MC). The primary goal of our study is to derive the precise geometry of binocular projections when the nodal point is placed at the anatomically correct location.

It should be noted that there is no single horopter. The shape and form of the horopter depends on its definition and the procedure used to measure it. We refer to Tyler (2004) for the historical and background information on binocular vision, including a comprehensive discussion of the many different notions of the horopter.

In particular, the empirical longitudinal horopter deviates from the V-MC. This so-called Hering-Hillebrand horopter deviation can be accounted for by asymmetry in the effective spatial positions of the corresponding elements in the two eyes. This deviation’s dependence on fixation distance is consistent with fixed corresponding points in retinal coordinates (Hillis & Banks, 2001).

The classical results of Helmholtz (1866) on the geometrical aspects of the horopter were recently extended by Schreiber and colleagues (2006). In particular, because of the extended Listing’s Law’s torsional adjustments at near distances, the horopter’s vertical component remains a straight line but with increasing slant with increases in fixation distance.

Here we study the geometric horopter based on the equality of visual angles between the two eyes. The visual angle between two external points is the angle subtended on the retina by the projecting rays toward those external points passing through the eye’s nodal point. We assume a simplified eye model with coincident visual and optical axes and an optical nodal point that is anterior to the eye’s rotation center. Further, we exclude cyclotorsion and assume that the eyes fixate on a single point in the visual plane. This plane contains the visual axes of the two eyes and is assumed, for simplicity, to be coplanar to the transverse (horizontal) visual plane of the head.

According to Müller (c. 1826), who was aware of Vieth’s geometrical work in binocular vision, the horizontal horopter is the circle through the fixation point and the nodal points of the two eyes, usually referred to as the V-MC. However, Müller assumed that the nodal point and the eye rotation center are coincident. An important implication of Müller’s assumption is well-known: When the fixation point changes along the V-MC, the vergence must be constant and the circle unchanged. Thus, the V-MC is the isovergence circle.

Gulick and Lawson (1976) realized that Müller’s assumption should be rejected because the anatomically correct locations of the nodal point and the rotation center are different. Their geometric analysis of the horopter used values averaged from several studies to locate the optical node of the primary visual axis to be at a point 6 mm anterior to the center of rotation and 17 mm anterior to the fovea. Figure 1 shows a schematic eye diagram with the nodal point N and the center of rotation C placed accordingly. Unfortunately, as we demonstrate here, their geometric study is flawed.

**Figure 1.**
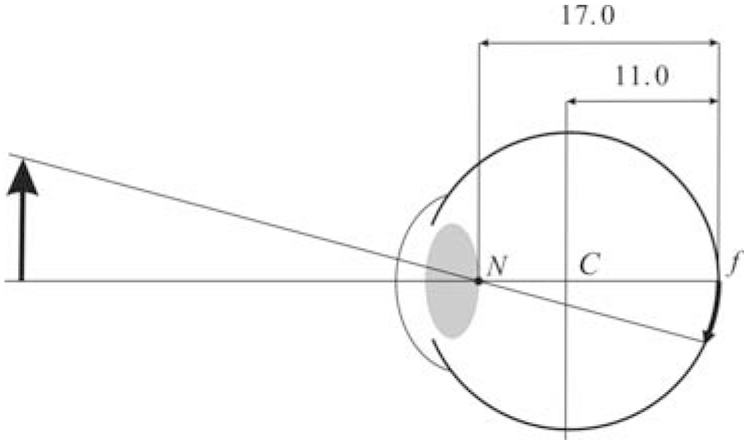
Eye model with the nodal point N and the eye rotation center C. The nodal point is located on the principal visual axis at a distance 17 mm anterior to the fovea f and the center of rotation is on the same axis at a distance 11 mm anterior to the fovea. These values are taken from Gulick and Lawson (1976).

We prove that, along the binocularly visible Vieth-Muüller circular arc connecting the eye rotation centers, there is an infinite family of horizontal horopters. They are parametrized by vergence, the specific fixation point along the corresponding isovergence (Vieth-Müller) arc, and the choice of the nodal point location. All the horizontal horopters for a constant vergence intersect at the point of symmetric convergence on the V-MC. Further, there is an infinite family of vertical horopters formed by straight lines parametrized by vergence. Each line is perpendicular to the visual plane and passes through the point of symmetric convergence. Thus, the complete picture is an infinite family of 3D geometric horopters (GHs) with two perpendicular components as described above. Further, we see that only the V-MC can be identified with the isovergence circle.

If we choose the location of the Cyclopean eye’s center to be on the shorter isovergence circular arc midway between the eyes’ rotation centers, then it follows that the fixation axis of the cyclopean eye is given by the version. Then, the Cyclopean visual axis’ rotation is simply the average of the left and right eyes’ rotations. Our choice of the Cyclopean eye location is motivated by this mathematical simplicity.

The main distinction between the GH and the V-MC is the relative disparity’s dependence on eye position when the nodal point has the anatomically correct location. We evaluate the difference between the GH and the V-MC for typical near fixation distances and discuss its significance for depth discrimination and other related functions of vision that make use of disparity processing.

## 2 The Horopter and Cyclopean Eye

In this section, we present a detailed geometric analysis of binocular projections. Table 1 gives a summary of the notation and important geometric facts used in this paper.

**Table 1.**
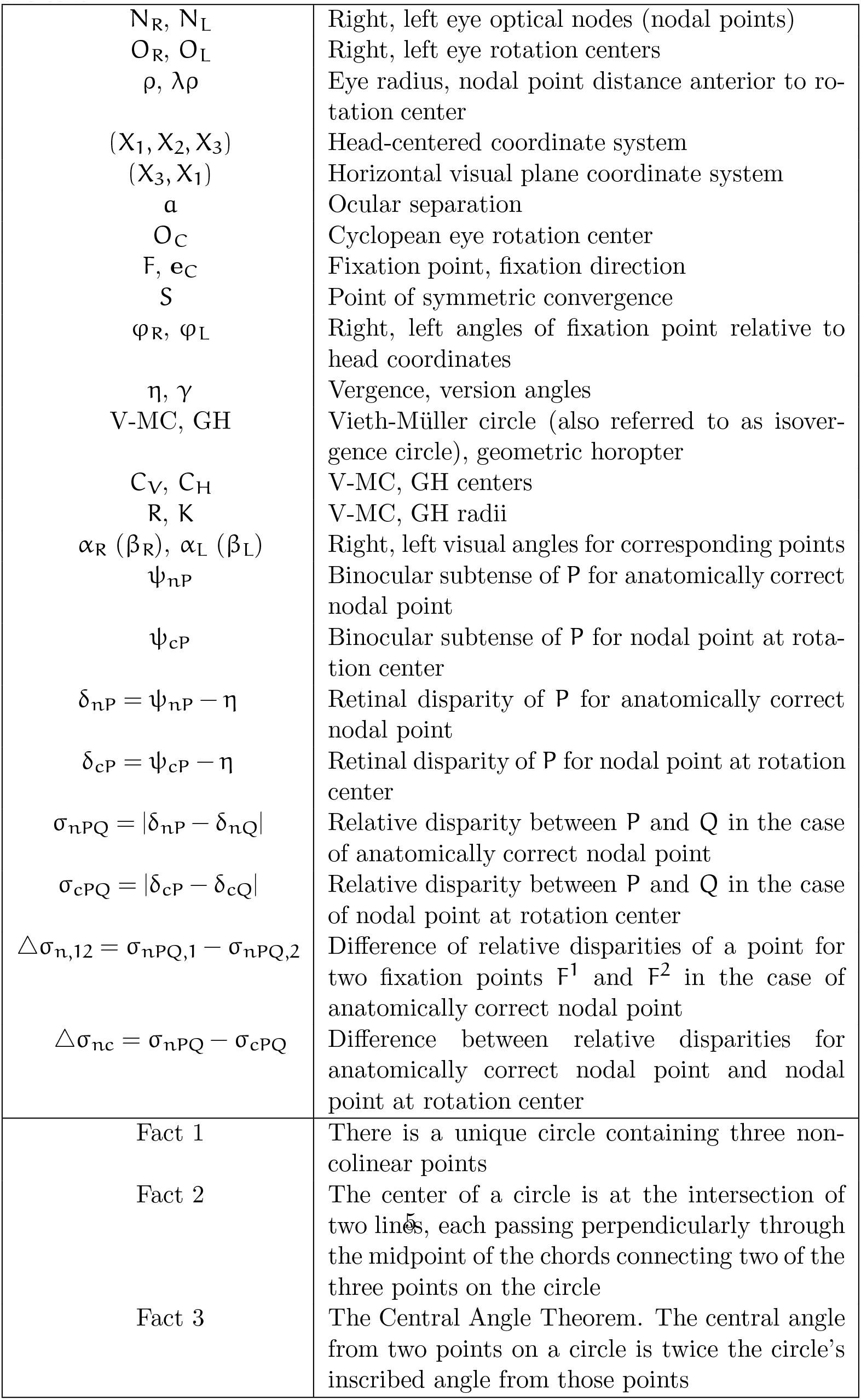
Summary of the notation and important geometric facts used in the text.

### 2.1 Geometric Formulation and Conjectures

We start with an introduction of the necessary notation and then discuss the definition of the GH in the visual plane.

The visual plane, according to our assumptions, contains the two eyes’ fixation points, nodal points, rotation centers and foveal centers. In Figure 2, the orientation of this plane is given by the right hand rule in the two-dimensional coordinate system (X_3_,X_1_) such that the head-centered coordinate system (X_1_, X_2_, X_3_) at origin O is positively oriented. For example, when the left and right eye fixate F, their positions are specified with respect to the head coordinates by φ_R_ > 0 and φ_L_ >0, respectively. Then, by definition, the vergence is η = φ_R_ – φ_L_.

**Figure 2.**
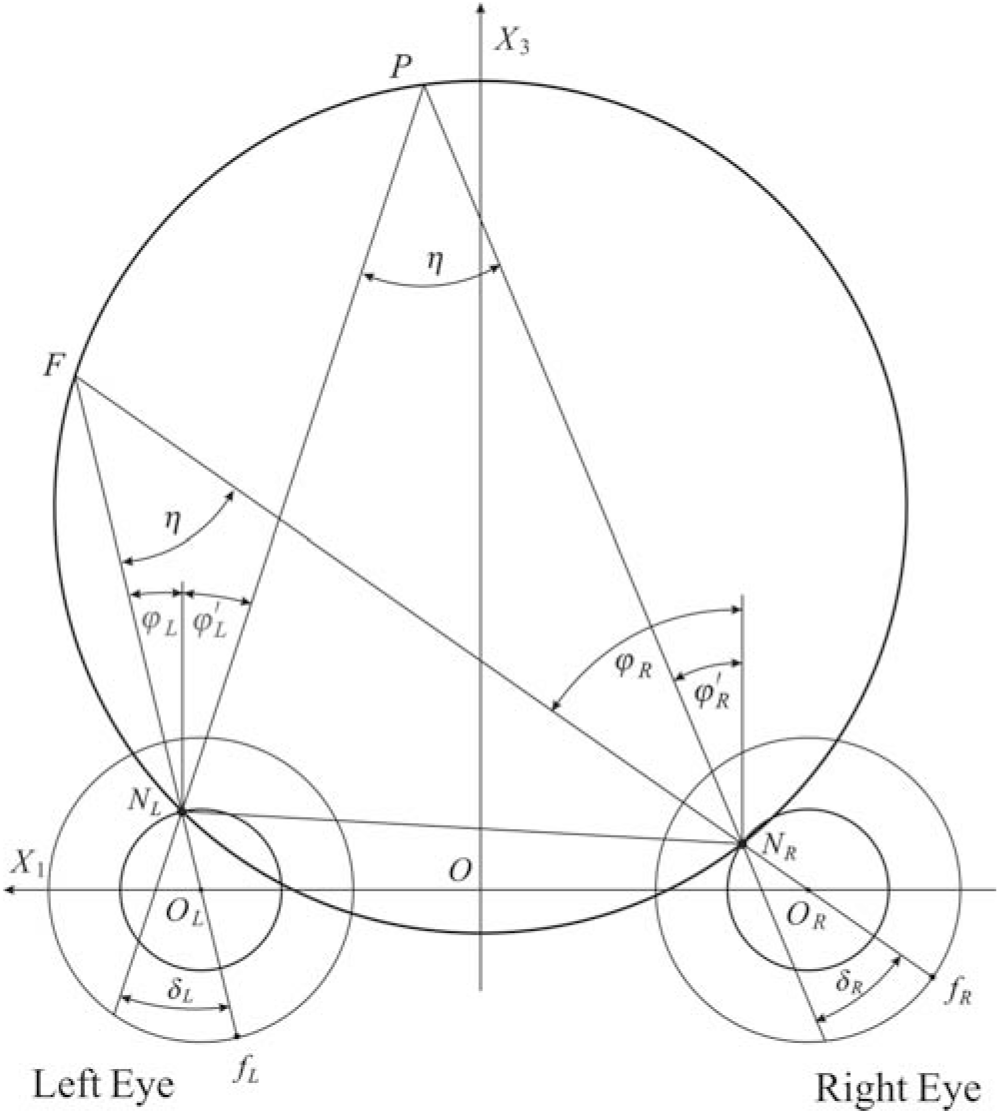
The GH is the curve containing the fixation point F and both the right and left nodal points N_R_ and N_L_. As explained in the text, all binocularly visible points on the curve subtend the same angle η relative to the two eyes’ nodal points (only one point P is shown).

The GH is the set of points in the visual plane subtending the same angle with the visual axes of both eyes. However, we explicitly demand that this definition involves the anatomically correct location of the nodal point, although later the proof is carried out for any nodal point located between the pupil center and the rotation center. As shown in Figure 2, the fixation point is F and the nodal points are N_R_ and N_L_ for the right and left eye, respectively.

We show now that if a point P is on the GH, then the disparity ∝_R_–∝_L_ = 0. The angles ∝_R_ and ∝_L_ are the visual angles of the point P in the right and left eye, respectively. It follows that the GH is formed by the circular arc that contains the point of fixation F and connects the nodal points N_R_ and N_L_. In fact, by the Central Angle Theorem, the angle at any point P on the circular arc is η. Therefore, η = φ_R_ – φ_L_ = φ′_R_ – φ′_L_. From the second equality we get φ_R_ – φ′_R_ = φ_L_ – φ′_L_. Taking into account the signs of the angles, this means that ∝_R_ = ∝_L_, i.e., the relative disparity is zero for all points on the GH as claimed.

Next, we explain the construction of the GH and V-MC for a given fixation point. The geometric facts used in the construction are the following. First, there is a unique circle containing three non-colinear points. These points in Figure 3 are F, N_R_ and N_L_. Second, the circle’s center is located at the intersection point of two different lines, both of which pass perpendicularly through the midpoint of the chord connecting two (of the above three) points. This condition also specifies the radius of the circle.

**Figure 3.**
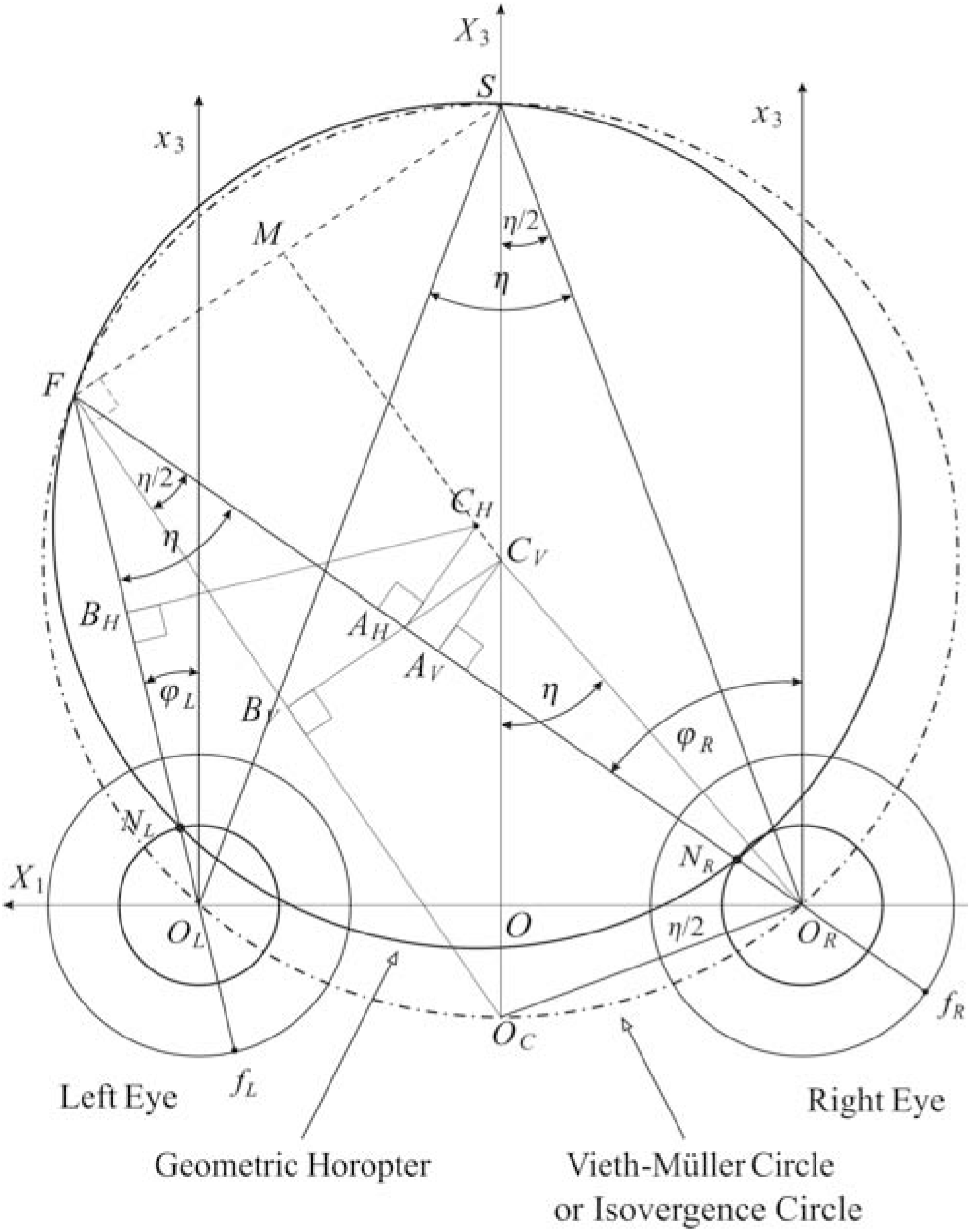
The construction of the GH and V-MC for the fixation point F in the visual plane. For the GH (solid line) two chords are FN_R_ and FN_L_. The perpendicular lines passing through the corresponding midpoints A_H_ and B_H_ intersect at the horopter center C_H_. A similar construction is used for the V-MC (dot-dashed line) with the center C_V_. When the eyes fixate on the points of the V-MC, the vergence remains constant. Conjectures inferred from the drawing are shown as dashed lines.

Further, by the Central Angle Theorem, we have the same angles η and η/2 at each of the vertices F and S. Finally, we note that the angle ∠O_C_O_R_O is equal to η/2 because ΔOO_R_O_C_ and ΔO_R_SO_C_ are similar triangles.

The dashed lines show the results suggested by the drawing. *Conjecture 1*: The point S, taken where the X_3_ axis intersects with the V-MC, lies on the geometric horopter as well. *Conjecture 2*: The line through points C_V_ and C_H_, the centers of the V-MC and the GH, is parallel to the line through Oc and F, and M is the midpoint of the line segment FS.

Obtaining conjectures from precise drawings is a nice feature of our approach and this is how we ‘discovered’ the correct geometry of binocular projection. Next, we will prove these conjectures.

### 2.2 Preliminary Results

In Figure 4, using the previously described method, we draw the GHs for the two fixations F and F′ on the V-MC (shown in green for F and brown for F′). It is enough to prove the conjectures for only one fixation point, say F.

**Figure 4.**
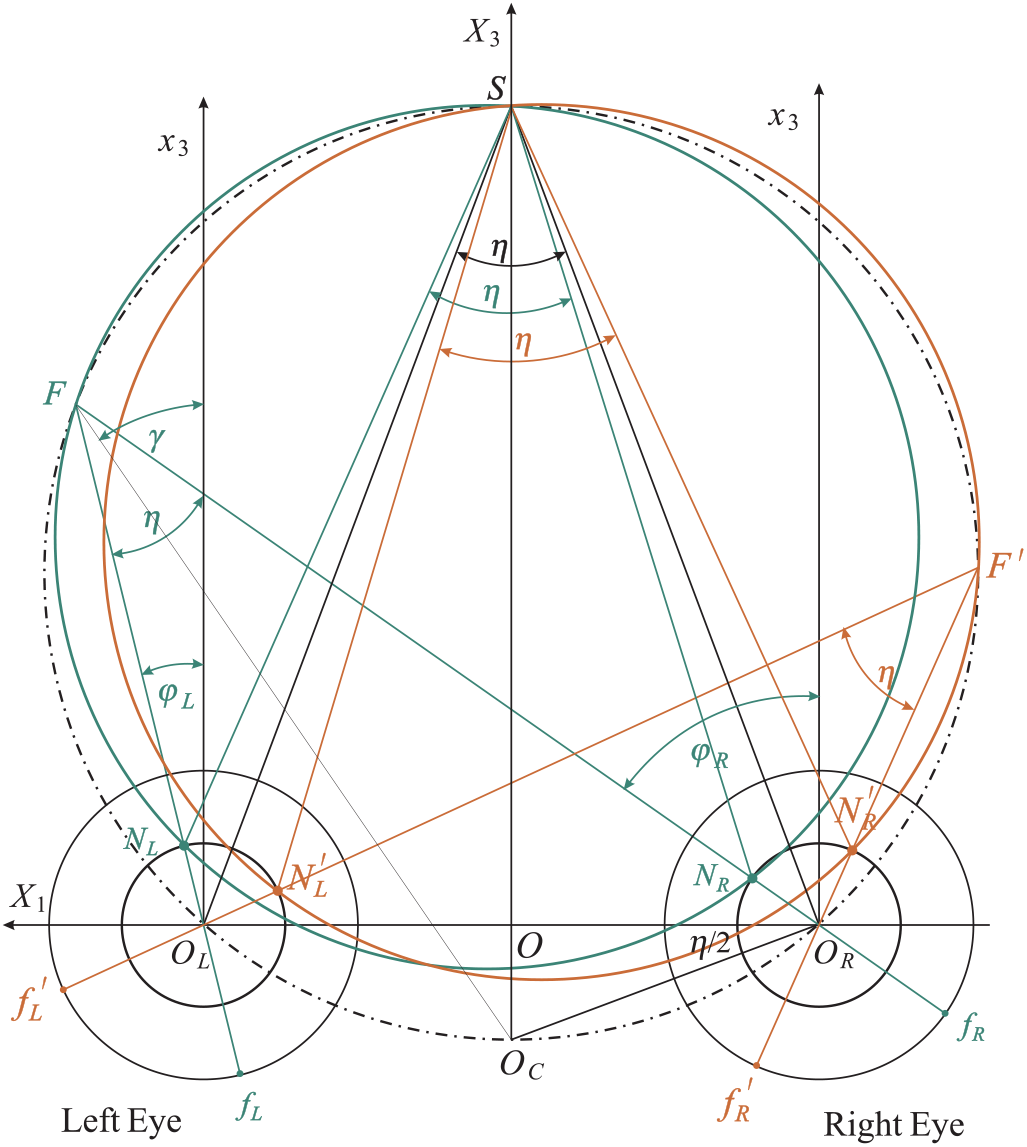
The GHs constructed by the previously described method are shown for the fixation points F and F′ on the V-MC. The point S of symmetric convergence is the point of intersection between the GHs and the V-MC. The proof of this fact is given in the text.

We first prove that, for all fixation points with a constant vergence, the corresponding GHs intersect at the symmetric convergence point S located on the V-MC. To prove this fact, we demonstrate that both the green triangle ΔSN_R_N_L_ and the triangle ΔSO_R_O_L_ have the angle η at the vertex S. To this end, it is enough to show that the triangles ΔSN_R_O_R_ and ΔSN_L_O_L_ are congruent. Since |O_R_S| = |O_L_S| and |O_L_N_L_| = |O_R_N_R_|, we only need to show that |N_R_S| = |N_L_S|.

Complex numbers significantly simplify proofs and are used whenever appropriate. To complete the proof, we introduce the following complex-number notation in (X_3_,X_1_) coordinates of the visual plane.

For the right eye,

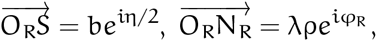

and for the left eye,

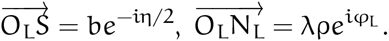

Then,

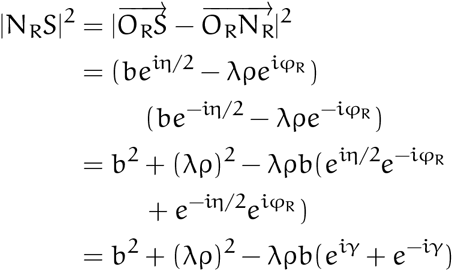

and, in a similar way,

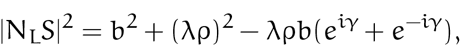

where we substituted version γ given by the expression

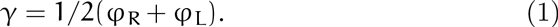

Thus, |N_R_S| = |N_L_S|, showing that ΔSN_R_O_R_ and ΔSN_L_O_L_ are congruent and hence that the vertex S must be on both the green horizontal horopter and the V-MC. This proves Conjecture 1.

The proof that was just completed clarifies two aspects of the GH. The first is that, in the visual plane, the locus of points lying on the horopter is the circular arc containing the fixation point and connecting the nodal points. It intersects the corresponding V-MC only at two points, the fixation point and the point of symmetric convergence. The second is that, elsewhere, the locus of these points is the line that is perpendicular to the visual plane and passes through the point of symmetric convergence. In particular, the GH is not the V-MC, unless one assumes an anatomically incorrect location for the nodal point, one that is coincident with the center of rotation.

### 2.3 Infinite Family of GHs

So far, the nodal point has been assumed to be located on the principal visual axis a distance λρ = 6 mm anterior to the eye rotation center. However, in our study of binocular geometry, we will assume an arbitrary 0 ≤ λ ≤ 1 to capture the way in which the GH depends on the choice of λ. Referring to Figure 1, the eye radius ρ and λ are well approximated by 11 mm and 6/11, respectively. Also, λ = 0 corresponds to Müller’s assumption that the nodal point is coincident with the eye rotation center and λ = 1 means that the nodal point is at the pupil center.

Figure 5 shows geometric details of the binocular projections for two nodal point locations, one at the anatomically correct location (λ = 6/11), and the other at the pupil center (λ = 1). It also shows our choice of position for the Cyclopean eye.

**Figure 5.**
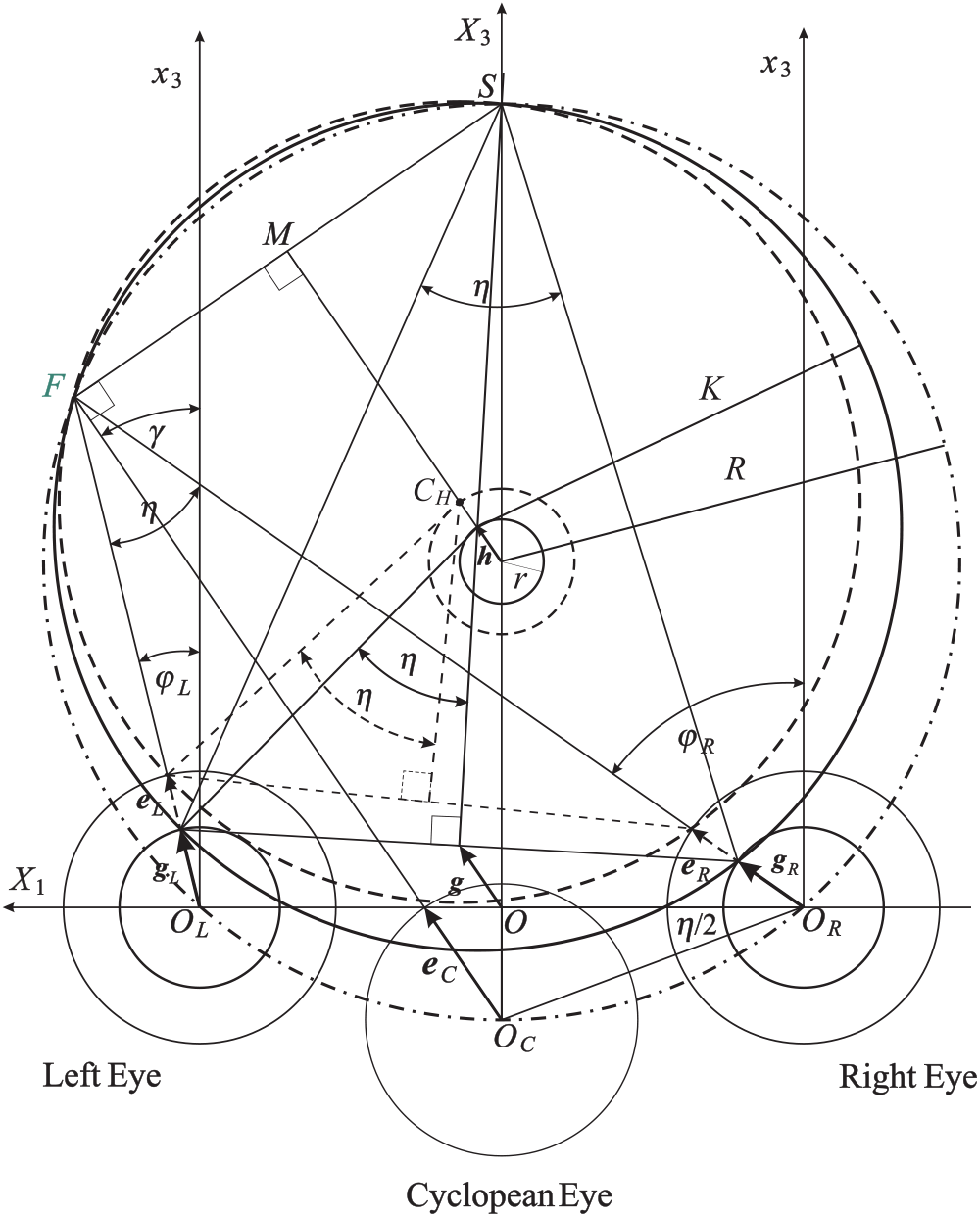
The detailed geometric description of the binocular projections. The GH for the anatomically correct nodal point location (solid line) is the circle of radius K and center C_H_. The horopter for which the nodal point is assumed to be at the location of the pupil center (λ = 1) is shown by the dashed lines. This horopter also intersects with the GH and the V-MC at the point S of symmetric convergence.

The geometric analysis we have described so far implies that the 3D GH consists of an infinite family with these two components. In the first component, the horopters in the horizontal visual plane are circular arcs containing the fixation point and connecting the nodal points. They are parametrized by the vergence, the specific fixation point within the V-MC, and the nodal point location (given by λ). For a fixed vergence value, all of the horizontal horopters intersect the corresponding V-MC at the point of symmetric convergence. The second component of the GR is the line that is perpendicular to the visual plane and passes through the point of symmetric convergence.

We now proceed to a detailed description of the GHs and the Cyclopean eye. Although our proof applies to all fixation points, we prove the results for only one fixation point. Further, to simplify expressions, we introduce the following vector notation (see Figure 5):

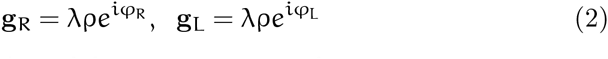

for the right and left eye’s nodal points, respectively.

We place the Cyclopean eye center on the V-MC at the intersection of the ray bisecting the vergence angles at the fixation points, and we take its orientation along the bisecting ray. It follows from the triangle ΔOO_R_O_C_ in Figure 5 that the Cyclopean eye center O_C_ is at a distance

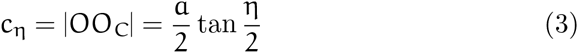

below the X_1_-axis passing through the eyes’ centers, where a is the ocular separation and η is the vergence. The visual axis of the Cyclopean eye for a fixation F is given by the version (Eq. 1).

According to the conventional theory, articulated originally by Hering (Hering, 1870) and augmented by others, the Cyclopean eye center is located on the interocular axis midway between the eyes. Then, a perceived direction, well-approximated by the version γ, is seen as if it were viewed from the Cyclopean position between the eyes (see, e.g., Banks, Van Ee & Backus, 1997).

Without resorting to discussion of ongoing theoretical and experimental studies on the many perceptual aspects related to the Cyclopean eye (including controversies, see for example Erkelens & van Ee, 2002; Ono, Mapp & Howard, 2002), its position as specified here is on the V-MC with the viewing direction precisely equal to the version for all choices of the nodal point location. This choice of the Cyclopean eye’s position is briefly discussed in the last section; but note that its position changes with changes in vergence.

To obtain the center of the horizontal horopter for the fixation point F, we choose one chord to be between fixation point F and point S on the X_3_-axis, and the other connecting the nodal points for arbitrarily chosen 0 ≤ λ ≤ 1. Because S and F are both on the green and dot-dashed circles, the line intersecting the centers of both circles is the line perpendicular to the line segment FS that passes through its midpoint M.

Thus, this perpendicular line contains the vector h between the centers. Further, FS is perpendicular to the bisecting ray at F that has the visual direction of the cyclopean eye given by the unit vector e_C_,

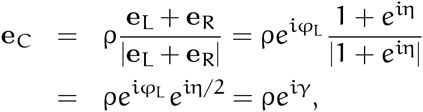

where we used the fact that vergence η = φ_R_, φ_L_, version γ = 1/2(φ_R_+φ_L_) and

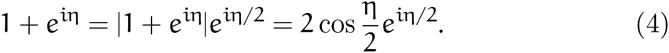

This proves that line segment O_C_F is parallel to C_V_C_H_, thus proving Conjecture 2.

Next, we demonstrate that, for a constant value of vergence, each horizontal horopter’s center lies on the circle whose center coincides with the center of the V-MC. First, we notice that the vector g, going from the head coordinates’ center O to the midpoint of the line segment connecting the nodal points, is the average vector

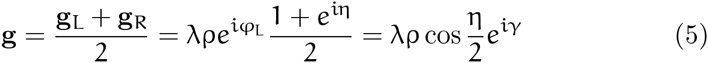

where we used (Eq. 4).

Thus, e_C_, g and h are parallel vectors. Next, using the fact that |OS| = 2R — c_η_, where R is the radius of the dot-dashed circle (V-MC) and c_η_ is given in Eq. 3, we conclude from the corresponding similar triangles (one with sides |OS| and |g| and the other with sides |C_V_S| and |h|), that the following proportion

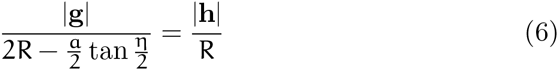

holds true.

Solving Eq. 6 for h and using Eq. 5, we obtain

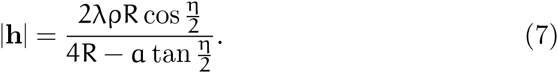

To simplify the last expression, we first obtain

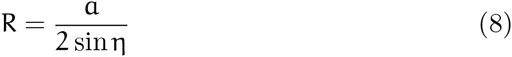

from the right triangle ΔOSO_R_ and Eq. 3.

Then, substituting Eq. 8 into Eq. 7, we derive

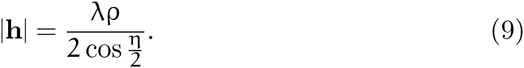

We see that h = |h|e^iγ^ and |h| is constant for fixed values of η and λ, i. e., it is independent of the version γ. Because Eq. 9 holds for all fixation points along the V-MC, we denote |h| by r so that

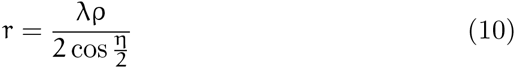

is the radius of the circle on which the centers of the GHs are located. Moreover, this circle has the same center as the V-MC.

Vergence decreases as fixation distance increases, so that Eq. 10 is well approximated by λρ/2 for large values of R. We take the fixation distance as 2R, which is justified for version values less than 15 deg.

### 2.4 The Center and Radius of the GH’s Circles

The GH center is the endpoint of the complex number

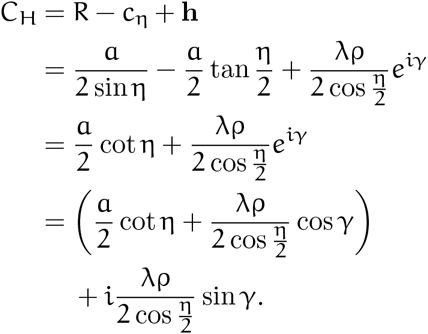

The radius K of the GH can be obtained from the identity

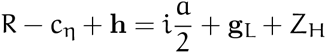

where ZH is the complex number representing the vector needed to obtain the equality. Then

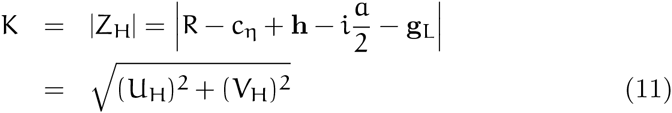

where

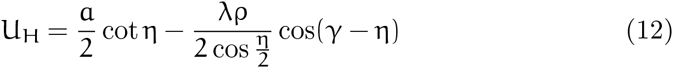

and

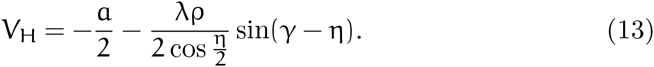

In the above expressions, we substituted Eqs. 2, 10, 8 and 3. Further, we used φ_L_ = γ — η/2 and the fact that h is parallel to the vector g (Eq. 5). We also have made use of standard trigonometric identities.

## 3 Numerical Study of the GHs

From Eqs. 8 and 10 in Section 2, we see that when the fixation distance (roughly in the range of 2R) increases, the vergence decreases to zero, while the radius r (Eq. 10) of the circle containing the centers of the GHs approaches a constant value λρ/2, or 3 mm for the anatomically correct GH.

In the remaining part of the paper, we assume the human average values of α = 65 mm, ρ = 11 mm and λ = 6/11. Using the expressions in Eqs. 8, 11 and 10, in Table 2 we show the values of R, K and r. These values are calculated for version values γ = 16°, 8°, 4° and vergence values of η = 6^o^, 8^o^, 10^o^.

**Table 2.**
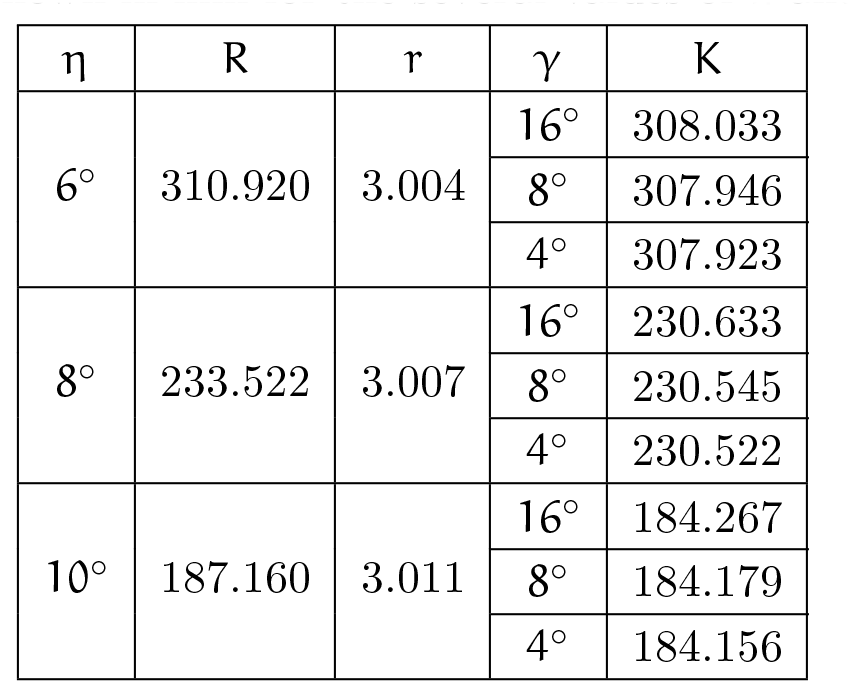
The values of R (Eq. 8), K (Eq. 11) and r (Eq. 10) are shown in mm for the several values of η and γ.

The fixation distance is defined here as the distance from the point O_C_ to the fixation point. We recall that, according to our choice for the Cyclopean eye position, the point O_C_ is the center of the Cyclopean eye and the visual axis’ direction is given by the version.

We estimate the fixation distance by 2R. From Table 2, the viewing distance is a typical near fixation for η ≤ 10° (Wong, Woods & Peli, 2002). We see that, for all viewing distances larger or equal to this near distance, the largest difference between the GH and the V-MC is about 3 mm near the point diagonally opposite to the fixation point. This visually irrelevant value is independent of the version.

Moreover, the largest distance between the GH and the V-MC that is visually relevant occurs midway between the fixation point and the point of symmetric convergence, both on the GH. This can be easily seen in Figure 5. Denoting this distance by ɛ_max_, we have from Figure 5

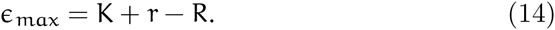

Because ɛ_max_ does not essentially change with the vergence when the version is kept constant, their values calculated for all cases in Table 2 are shown in Table 3 only for version values of 16°, 8° and 4°.

**Table 3.** The distance ɛ_max_ (Eq. 14) in mm for the cases shown in Table 2.

| $\gamma$ | $16^\circ$ | $8^\circ$ | $4^\circ$ |
| --- | --- | --- | --- |
| $\epsilon_{\max}$ | 0.118 | 0.030 | 0.007 |

The values of ɛ_max_ in Table 3 are quite small. Their visual significance is discussed in Section 6.

## 4 Relation to Previous Literature

Gulick and Lawson (1976) attempted to determine the effect the separation of the center of rotation from the optical nodal point has on binocular geometry. In Chapter 3 the authors first discuss Müller’s horopter (V-MC) and the later modification by Graham (1965) that placed the coincident optical nodal point and rotation center anterior to the geometrical center of the eye. Because of these anatomically incorrect assumptions, they rejected both models and presented their binocular geometry under the assumption that the nodal point is 6 mm anterior to the eye’s center of rotation, and that the rotation center coincides with the eye’s geometric center (Figure 1).

However, with reference to our Figure 5, they erroneously concluded that F falls on the horopter of S, but S does not fall on the horopter of F [see page 81 and Figure 3.10 in Gulick and Lawson (1976)]. According to our geometric analysis (Figure 5), the correct statement should be that S falls on the horopter of F, but F does not fall on the horopter of S. This applies to all fixation points on the V-MC such that F ≠ S. Gulick and Lawson (1976) go on to conclude that the GH is irrelevant for a perceptual analysis of binocular vision. However, we have shown that with a correct geometric analysis of the GH, retinal disparities depend on fixation within the V-MC, a fact that is highly relevant for understanding the perceptual interpretation of disparity.

## 5 Retinal and Relative Disparities

Binocular disparity (or stereopsis) refers to the small differences in the perspective projections on the right and left eyes that result from the eyes’ lateral separation. When a point lies in front or behind the horopter curve containing the fixation point, the difference in the angles subtended on each retina between the image and the center of the fovea defines absolute retinal disparity. This difference provides a cue for the depth of the object from an observer’s current point of fixation. The difference of retinal disparities for a pair of points defines their relative disparity. The relative disparity provides a cue for the perception of 3D structure such as relative depth and shape. It is usually stated that relative disparity does not depend on the eye position (Marr, 1985).

Because binocular geometry is different for the anatomically correct GH and for the V-MC, we need to investigate whether the relative disparity is indeed independent of the eyes’ positions. In what follows, we use the subscript ‘n’ for quantities obtained using the anatomically correct location of the nodal point at λρ = 6 anterior to the center of rotation, and the subscript ‘c’ for quantities obtained when the nodal point coincides with the eye rotation center (λ = 0).

The sign of visual angle conforms to the orientation of the fixation plane’s coordinate system (X_3_,X_1_) discussed before. For example, in Figure 6, the visual angle of projections ∝_nR_ and ∝_nL_ for the endpoint P are negative while the projection angles ß_nR_ and ß_nL_ for the endpoint Q are positive.

**Figure 6.**
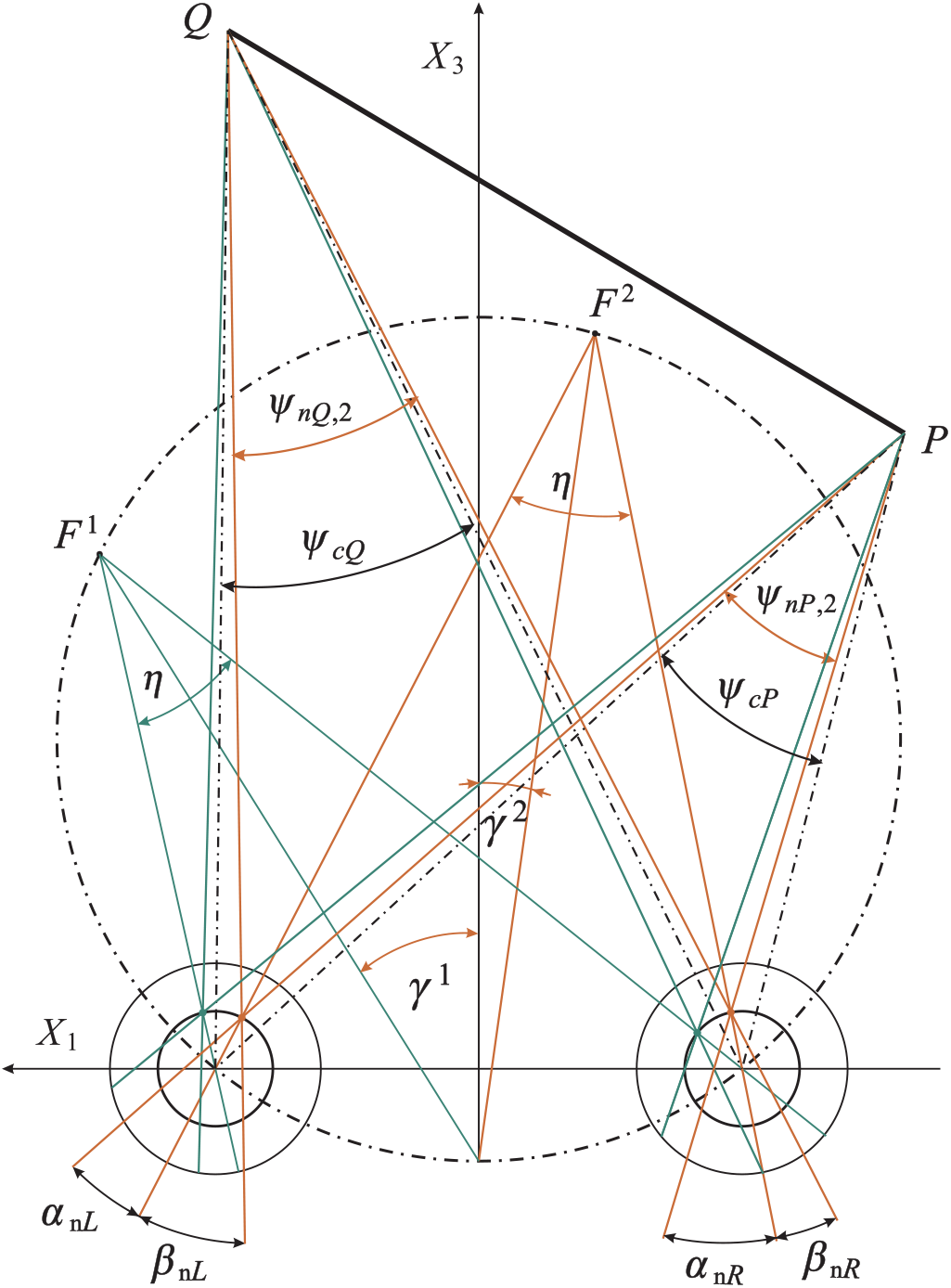
The drawing used to define and calculate disparities in terms of the binocular subtense. The subtense of the points P and Q are the angles ψ_np,2_ and ψ_nQ,2_ for the case of λ = 6/11, and ψ_c_p and ψ_c_Q for the case of λ = 0. The subtense angles are shown only for the fixation F^2^. We note that the binocular subtense is independent of the eye’s position for the case λ = 0.

By definition, the retinal disparities of endpoints P and Q in the case of the anatomical position of the nodal point (subscript *n*) are:

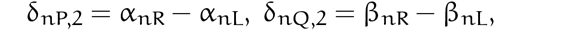

where the visual angles α of the endpoint P and the visual angles β for Q, each angle for the right eye (subscript R) and for left eye (subscript L), are shown in Figure 6.

From Figure 6 one can obtain the expressions for the retinal disparities δ_np,2_ and δ_nQ,2_ and the relative disparity σ_nPQ,2_ in terms of the endpoints’ binocular subtense ψ_np,2_ and ψ_nQ,2_ (see also page 37 in Howard and Rogers (1995)), as follows:

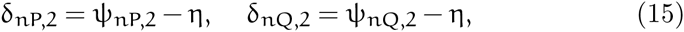

and

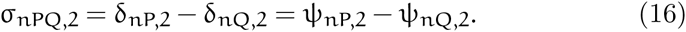

However, because δnp,2 and δ_nQ,2_ in Eq. 15 contain information about the fixation point but σ_npQ,2_ in Eq. 16 does not, we choose relative disparity for the line segment with the absolute value

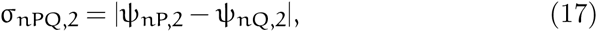

avoiding reference to any retinal landmarks. This formula does not account for the line segment orientation.

The formulas in Eqs. 15 and 17 are especially convenient because they are easy to calculate either from the drawing or from geometric formulas when the coordinates of the points are known.

For the subscript c (i.e., for λ = 0),

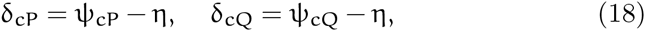

and

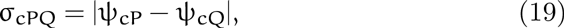

where ψ_c_p and ψ_c_Q are the angles between the dot-dashed lines at P and Q, shown in Figure 6. We note that relative disparity (Eq. 19) is independent of the eye’s position.

In Figure 7, we show the geometric parameters of the binocular projections. Further, 2R/ρ = 25.3, where ρ is the eyeball radius and R is the radius of the V-MC. Also, the ratio α/ρ = 6.0, where a is the interocular distance. These values mean that if we scale the drawing such that ρ = 11 mm, the distance to fixation points F^1^ and F^2^ is about 2R = 28 cm, and α = 6.6 cm. Also, in this case, the distance to the fixation point F^3^ is 23 cm.

**Figure 7.**
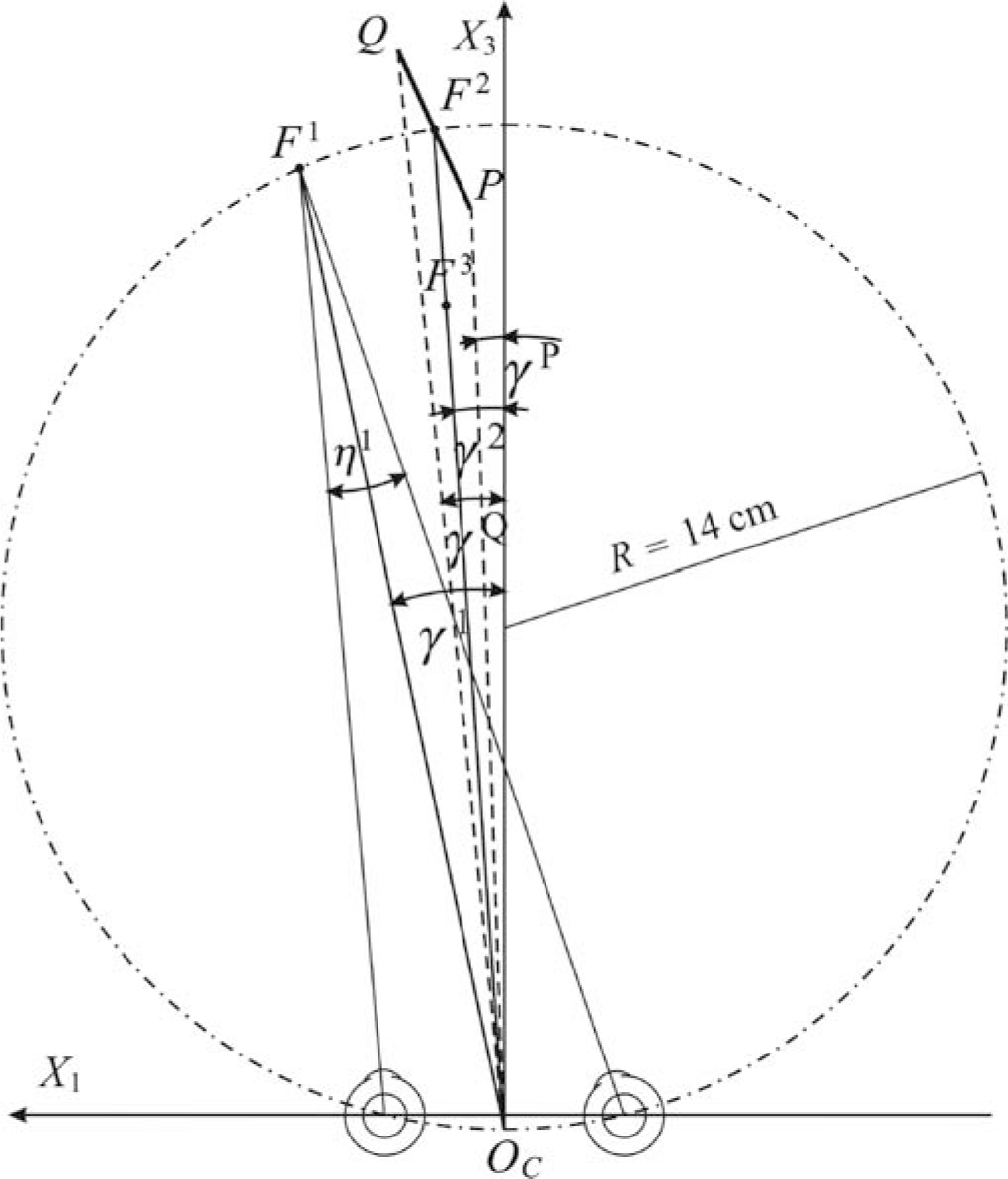
The binocular projection parameters for which the retinal and relative disparities are calculated. Here, the version values for the fixation points are γ^1^ = 12°. and γ^2^ = 4°., and the corresponding angles of the endpoints are γ^P^ = 2°. and γ^Q^ = 5.36′. The vergence for the fixation points F^1^ and F^2^ is η^1^ = η^2^ = 13.45′11˝ and the vergence for the fixation point F^3^ is 16°.42′47˝. Also, the distance from O_C_ to the fixation points F^1^ and F^2^ is about 28 cm and the distance to the fixation point F^3^ is 23 cm.

The retinal disparities in Figure 7 can be calculated using data from Table 4 and Eqs. 15 and 18.

**Table 4.** Binocular subtense of P and Q for the fixation points F^1^, F^2^ and F^3^ calculated for parameters in Figure 7.

| Model | Vergence | Point P | Point Q |
| --- | --- | --- | --- |
| V-MC | 13°45'11" | $\psi_{cP}$ | $\psi_{cQ}$ |
|  |  | 15°00'36" | 12°39'58" |
| GH | 13°45'11" | $\psi_{nP,1}$ | $\psi_{nQ,1}$ |
|  |  | 15°02'42" | 12°38'53" |
| | | $\psi_{nP,2}$ | $\psi_{nQ,2}$ |
|  |  | 15°01'55" | 12°38'42" |
| | 16°42'47" | $\psi_{nP,3}$ | $\psi_{nQ,3}$ |
|  |  | 14°57'58" | 12°35'20" |

The relative disparities σ_npQ,1_, σ_npQ,2_, σ_cPQ_, and their differences

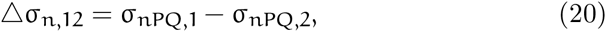

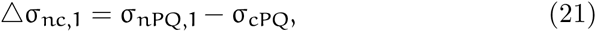

and

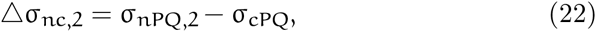

derived using the data in Table 4 are shown in Table 5.e

**Table 5.** Relative disparities for the endpoints of line segment PQ and their differences (see text for definitions).

| Relative Disparity |  |  |  |
| --- | --- | --- | --- |
| $\sigma_{nPQ,1}$ | $\sigma_{nPQ,2}$ | $\sigma_{nPQ,3}$ | $\sigma_{cPQ}$ |
| 2°23'49" | 2°23'13" | 2°22'38" | 2°20'38" |
| Relative Disparities Difference |  |  |  |
| $\Delta\sigma_{n,12}$ | $\Delta\sigma_{n,23}$ | $\Delta\sigma_{nc,1}$ | $\Delta\sigma_{nc,2}$ |
| 36" | 35" | 3'11" | 2'35" |

From the geometry that arises when the nodal point is placed at its anatomical location, we see that relative disparity depends on eye position.

The significance of the dependence of relative disparity on the fixation point will be discussed in the next sections for retinal disparity calculated for the fixation distance about 270 cm (for the radius R = 134.7 of the corresponding V-MC) such that the vergence at F^1^ and F^2^ is 1°22′55˝. The object PQ is the same as in Figure 7 except that it is translated along the line into the new position F^2^ (on the V-MC of radius 134.7 cm) so that the version angles (γ) shown in Figure 7 are preserved. The corresponding retinal disparities are shown in Table 6.

**Table 6.** Retinal disparities of P and Q for the fixation points F^1^ and F^2^ as in Figure 7 but calculated for the fixation distance of 270 cm

| Model | Vergence | Point P | Point Q |
| --- | --- | --- | --- |
| GH | $1^\circ 22' 55''$ | $\delta_{nP,1}$ | $\delta_{nQ,1}$ |
| | | $14' 46''$ | $-2' 09''$ |
| | | $\delta_{nP,2}$ | $\delta_{nQ,2}$ |
| | | $14' 28''$ | $-1' 48''$ |

The relative disparities σ_nPQ,k_, k = 1,2 and their difference Δσ_n,12_ are the following:

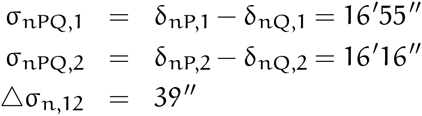

## 6 Discussion

Conventional theory of binocular projection based on the V-MC incorrectly assumes the eye’s nodal point and rotation center share the same location. The precise, but simple, binocular projection geometry presented here corrects this conventional theory. It is well known that, when the eyes fixate on the points of the V-MC, the vergence and the circle remain unchanged. This property is not shared by the anatomically correct GH.

Our geometric analysis showed that, when the eyes fixate a point on the binocularly visible arc of the V-MC in the horizontal visual plane, there is an infinite family of GHs. The horizontal horopters are formed by circular arcs connecting the nodal points of the two eyes and they intersect at the point of symmetric convergence. Apart from that, for each constant vergence, there is a vertical horopter consisting of a straight line that is perpendicular to the visual plane and passes through the point of symmetric convergence.

Thus, the complete picture of the GH involves an infinite family of 3D GHs with two perpendicular components, as described above. Although our geometric analysis has been made with the assumption that the visual plane is horizontal, the same geometry applies when the fixation target is elevated or depressed, as long as one ignores the typical cyclotorsion that occurs outside the horizontal plane in binocular vision.

Further, we chose the position of the Cyclopean eye by specifying its center’s location to be on the shorter isovergence circular arc that is midway between the eyes’ rotation centers. Only for this choice does the Cyclopean eye’s fixation axis’ direction correspond to the version. This position of the Cyclopean eye has a particularly simple property: its rotation is the average of the two eyes’ rotations. We note that if the Cyclopean eye’s location is assumed to be on the interocular line midway between the two eyes, the visual axis’ rotation would be a complicated function of the left and right eyes’ rotations. This simple properties of the Cyclopean eye position is important in our modeling of binocular vision for anthropomorphic robots (Turski, 2010).

We carried out two numerical studies. First, we investigated the question of how well the GH is approximated by the V-MC for typical near viewing distances. Second, we demonstrated an important consequence of the anatomically correct binocular projections: relative disparity dependends on the fixation point.

However, to understand the impact these results could have on human binocular vision, we need to discuss stereopsis and other visual functions that make use of disparity processing.

We normally do not experience double vision for points in a narrow band around the horopter, known as Panum’s fusional area. Originally, this area was proposed by Panum in 1858 for the empirical horopter (see, e.g., Verhoeff, 1959) and is used here for V-MCs and GHs. Experiments with empirical horopters found the width of this region to be about 0.5 deg in the vicinity of the fovea (Ponce & Born, 2008) and slightly increasing at larger eccentricities. However, some authors give smaller values. If an object falls within this region, it will be seen as a single object that is offset in depth relative to the fixation point. As disparity increases, fusion will be lost at some point.

Nevertheless, the visual system can extract meaningful depth information for disparities up to several degrees, depending on stimulus size, spatial and temporal frequency (see Ponce & Born, 2008). This could be important for minimizing bothersome double vision when objects are projected into the macula, the retinal region that extends about a few degrees from the foveola center, the region that is responsible for detailed central vision.

The spatial range of stereopsis is set by the minimum retinal disparity that can be resolved. The stereoacuity threshold under ideal conditions (high contrast, sharp edges and viewing at about 40 cm) is in the range of 2-6 arc sec (or 0.004-0.01 mm) (Wilcox & Harris, 2010). Also, the best stereoacuity is about 0.25 degrees (or 0.07 mm) from the foveola center.

How do these data compare with our analysis of the GH approximation by V-MC? Using the results in Table 3, we see that even for a version of 4°, the maximum value of visually relevant distance between the GH and V-MC of is about 0.007 mm and occurs at the horopter’s midpoint between 0° (the point of symmetric convergence) and 4° (the fixation point). Thus, a point on the V-MC will have zero depth relative to another nearby point on the V-MC, but they could be seen as offset in depth relative to the GH and maybe even relative to each other.

Given that the spacing between cones in the fovea is on the order of 30 arc sec, human discrimination of 5 arc sec of retinal disparity is extraordinary and termed a hyperacuity. Visual computations that make use of fine-scale binocular disparity information include ‘breaking camouflage’ when the outline of an object with pattern matching surroundings ‘pops out’ from the background because the object and the background are at slightly different depth. This figure-ground segmentation’s use of disparity processing is vividly demonstrated in the ‘Magic Eye’ images invented by Christopher Tyler in 1979, a postdoc of Bela Julesz.

A commonly held belief is that relative disparity is invariant under the change of the eyes’ positions. However, this is true only in conventional binocular theory, which is based on the V-MC. We show in Table 6 a dependence of relative disparity on the eyes’ fixation. In particular, Table 6 shows calculations for typical viewing parameters and suggests changes in disparity that could have a noticeable impact on vision. They are well within binocular acuity limits. We hypothesize that such disparity changes could lead in changes of perceived object shape with changes in fixation (across a saccade). This is an important observation because the eyes reposition gaze in natural viewing with, on average, 4 saccades per second. Even during fixation the eyes continually jitter, drift, and make micro-saccades. It was proposed by Vlaskamp et al. (2011) that these tiny eye movements during fixation cause changes in disparity estimation that are similar to spatial blur.

However, we wish to hypothesize that the small changes in perceived size and shape due to eye movements may be needed, not only for the perceptual benefits such as ‘breaking camouflage’, but also for the aesthetic benefit of stereopsis. For example, this can be demonstrated by the following quotations. Formerly stereoblind adults, when they became aware of the three-dimensionality of the visual world for the first time, are quoted by Ponce and Born (2008). One person wrote, ‘Before my vision changed I would not have said that the tree looked flat, but I had no idea just how round a tree’s canopy really is … When I began to see with two eyes, everything looked crisper and much better outlined.’ Another person wrote, ‘Everything has edges!’

Of course, the suggested impact of the dependence of relative disparity on the fixation point can only be confirmed by experiments. Since stereopsis’ functional significance has been rather neglected from the time it was first explained by Charles Wheatstone in 1838 (see Fielder & Moseley, 1996), such experiments are worth undertaking.

## Acknowledgment

We thank the reviewers for their comments and for pointing out the Gulick and Lawson (1976) reference. We also acknowledge editorial help of Michael Landy. Special thanks to Alice Turski, an MD/PhD student at NYU, for her many long discussions that improved the presentation.

## Footnotes

To appear in Vision Research

1 Retired

